# Inflammatory mediators act at renal pericytes to elicit contraction of vasa recta and reduce pericyte density along this medullary vascular network

**DOI:** 10.1101/2022.06.21.496998

**Authors:** Rebecca J. Lilley, Kirsti Taylor, Scott S.P. Wildman, Claire M. Peppiatt-Wildman

**Affiliations:** University of Kent; Northeastern University

## Abstract

Regardless of initiating cause, renal injury promotes a potent pro-inflammatory environment in the outer medulla and a concomitant sustained decrease in medullary blood flow (MBF). This decline in MBF is believed to one of the critical events in the pathogenesis of acute kidney injury (AKI), yet the precise cellular mechanism underlying this are still to be fully elucidated. MBF is regulated by contractile pericyte cells that reside on the descending vasa recta (DVR) capillaries, which are the primary source of blood flow to the medulla. Using the rodent and murine live kidney slice models, we sought to investigate the acute effects of key medullary inflammatory mediators TNF-α, IL-1β, IL-33, IL-18, C3a and C5a on vasa recta pericytes. Live kidney slices taken from both mice and rats and exposed to TNF-α, IL-18, IL-33, and C5a demonstrated a real-time pericyte-mediated constriction of DVR. When pro-inflammatory mediators were applied in the presence of the AT_1_-R blocker Losartan the inflammatory-mediated constriction that had previously been observed was significantly attenuated. When live kidney slices were exposed to inflammatory mediators for 4-hours, we noted a significantly reduction in the number of NG2+ positive pericytes along vasa recta capillaries in both rodent and murine kidney slices. Data collected in this study, demonstrate that inflammatory mediators can dysregulate pericytes to constrict DVR diameter and reduce the density of pericytes along vasa recta vessels, further diminishing the regulatory capacity of the capillary network. We postulate that preliminary findings here suggest pericytes play a role in AKI.

**New & Noteworthy:** How medullary blood flow (MBF) becomes disproportionately dysregulated following renal injury is poorly understood yet is associated with worse prognostic outcomes following AKI. This study shows in both rats and mice that inflammatory mediators associated with AKI have acute and sustained microvascular actions at pericytes eliciting dysregulation of descending vasa recta (DVR) diameter and their loss from the DVR. This work highlights a possible pathology behind the dysregulation and reduction of MBF observed following AKI.

## Introduction

Acute kidney injury (AKI) is a global health concern, with ∼13 million cases and ∼1.4 million deaths per year (1). Regardless of the instigating injury, renal diseases have inflammation as a common underlying pathogenic mechanism (2). Rapidly post-injury, inflammation is initiated with infiltrating immune cells (3,4) and high levels of pro-inflammatory mediators propagating further inflammation and damage. Haemodynamic alterations are also an early notable feature in renal injury(4). Importantly, most evidence indicates the renal region primarily affected is the medulla with a disproportionate dysregulation of blood flow (MBF) (5)(Freitas and Attwell, 2021). In animal models of ischemia reperfusion (IRI), despite post-reperfusion recovery of cortical blood flow, after a transient improvement post-reperfusion in the medulla, MBF follows a progressive gradual decline of up to 50% (6). Further still, this dysregulation of MBF is thought to be a critical event in the pathogenesis of AKI to CKD, yet the mechanisms behind this remain unclear. Blood flow to the medulla is provided by the vasa recta capillaries (7). Interestingly, the infiltrating cytokines and complement proteins that residential immune cells secrete, are both directly and indirectly vasoactive (8), with the involvement of Angiotensin-II (9). Whilst the inhibition of TNF-α, IL-18, IL-1β and C5a, amongst other inflammatory mediators, has shown to reduce inflammatory injury and preserve renal function (4,10), how these inflammatory mediators may be implicated in the dysregulation of renal blood flow, specifically in the sustained dysregulation of MBF in AKI is not well characterised.

Our laboratory (11,12), and others (7,13), have demonstrated that the cellular regulators of MBF are resident contractile pericytes, which respond to vasoactive cues from neighbouring endothelial and tubular cells to regulate vasa recta diameter and thus in turn alter flow through these capillaries (12,13). Pericytes have been implicated in progressive chronic kidney disease (CKD), with genetic fate-mapping studies demonstrating pericytes are the major source of myofibroblasts following injury (14–17), and their detachment and loss underpins the microvascular rarefaction associated with the progression of AKI to CKD (14,18). Recent work has further highlighted the role of vasa recta pericytes in the medullary no-reflow phenomenon post-reperfusion following renal ischaemia (19). The response of pericytes to the early onset renal inflammation that underpins AKI remains less clear but the delineation of underlying signalling mechanisms driving this inflammatory response may support our identification of novel targets that are involved in the resultant dysregulation of pericyte-mediated changes in vasa recta diameter, and by extension MBF. As such, the aim of the present study was to use both rodent and murine live kidney slice models, in combination with imaging techniques, to investigate the effect of inflammatory mediators on renal pericyte-mediated regulation of vasa recta capillaries.

## Materials and Methods

### Tissue Slicing

Animal experiments were conducted in accordance with United Kingdom Home Office Scientific Procedures Act (1986). Adult male Sprague-Dawley rats (200-225 g) or adult male C57BL/6J mice (63-70 days; purchased from Charles river UK Ltd, Kent, UK) were killed by cervical dislocation and kidney tissue slices were obtained as previously described (11).

### Live tissue DIC imaging experiments and Analysis

Live tissue DIC imaging experiments were performed using a method, previously described (Crawford et al., 2011, 2012). Video images of live tissue were collected to determine the effect of the inflammatory mediators on vessel diameter. Kidney slices were superfused with PSS alone (baseline), followed by PSS containing an inflammatory mediator, and then subsequently subjected to a PSS wash. Time-series analysis of kidney slice experiments was carried out off-line using the public domain software ImageJ (NIH, http://rsb.info.nih.gov.ij), as previously described (11).

### Anti-NG2 immunohistochemistry and image analysis

Immunohistochemical experiments with kidney slices were used to investigate the effect of pro-inflammatory mediators on renal pericyte morphology and density. These were performed as previously described (Crawford et al., 2012). Live tissue was exposed to inflammatory mediators for 4-hours prior to fixation. Alexa fluor 488-conjugated isolectin B_4_ (IB_4_; I21411, Invitrogen Ltd.) was used to identify the vasa recta capillaries and anti-neural-glial 2 (NG2; AB5320, Merck-Millipore) was used as a marker of pericytes. Anti-NG2 primary antibody was probed with donkey anti rabbit Alexa 555 (A-21208, Invitrogen Ltd.) secondary antibody. Images were analysed off-line using ImageJ (NIH, http://rsb.info.nih.gov.ij).

### Statistical Analysis

For analysis of the effect of inflammatory mediators (*treatment*) on pericyte density *(density*), R packages *lme4*(https://cran.r-project.org/web/packages/lme4/index.html) and *emmeans (https://cran.rstudio.com/web/packages/emmeans/)* were used. Given the overdispersed nature of our count data, a negative binomial regression was used, with the model having the form *density ∼ treatment + (1* | *animal)*. Assigning animal as a random effect was to account for the technical replicates in the data and within animal variation. We checked for violations of model fit for this model using a QQ-plot from the *DHARMa package* (https://cran.rstudio.com/web/packages/DHARMa/index.html) and a fitted versus Pearson residuals plot.

Graphpad PRISM 5.0 was used for statistical analysis of all normally distributed data sets. For DIC experiments, and murine NG2 experiments, statistical significance was calculated using a two-tailed Student’s *t*-test that was paired or unpaired when relevant; P < 0.05 was considered significant. Regarding all other experiments, statistical significance was calculated using one-way ANOVA and post hoc Dunnett test; P < 0.05 was considered significant. All values presented are expressed as mean ± SEM; number of animals (n).

## Results

### Inflammatory mediators evoke pericyte-mediated changes in in situ vasa recta capillary diameter

Fig. 1A shows a typical field of view of a vasa recta capillary (DVR; Ai-ii) and corresponding temporal response profile (Aiii) to an agonist. To determine the acute effects of exposure of rat medullary DVR capillaries to key innate immune components, live kidney slices were superfused with C5a, IL-33, TNF-α and IL-18 (10 ng/ml); which resulted in a pericyte-mediated constriction of vasa recta capillaries. In all cases constriction of capillaries was significantly greater at pericyte sites (15.2%±1.2%, 9.2%±0.8%, 11.3% ± 1.4%, 9.8% ± 0.7%, respectively; Fig. 1B) than at non-pericyte sites (1.0% ± 0.5%; n = 12; 1.9% ± 0.6%; n = 7; 0.6% ± 0.3%; n = 9; 0.7% ± 0.5%; n = 5; n = 3; respectively; ***P < 0.001, Fig. 1B). The C5a-, IL-33-, TNF-α- and IL-18-mediated constriction of DVR was reversible at pericyte sites in 52%, 37%, 38% and 47% of experiments performed respectively. Exposure of live tissue to C3a (10 ng/ml) and IL-1β (10 ng/ml) failed to elicit a significant greater change in vasa recta capillary diameter at pericyte sites (3.4% ± 0.5%, 2.6% ± 0.8%; data not shown) compared to non-pericyte sites (2.0% ± 0.8%; n = 5; 0.6 % ± 0.2%, n = 10; P > 0.05; data not shown).

**Figure 1.**
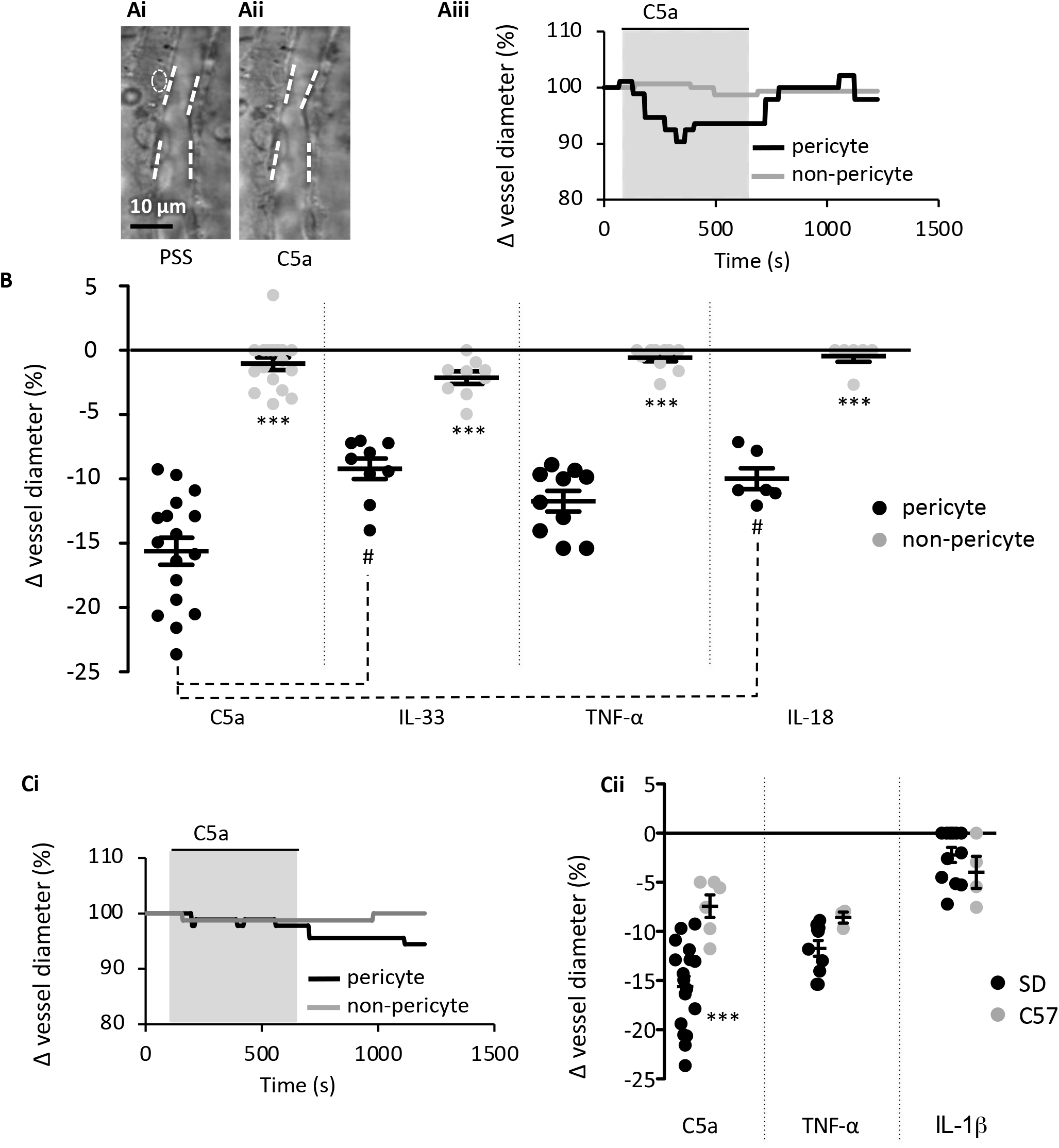
Innate immune components evoke a pericyte mediated constriction of rodent and murine vasa recta. Images show a typical field of view of Sprague-Dawley (SD) *in situ* vasa recta during superfusion of tissue with PSS (Ai) and C5a (Aii). Pericytes can be seen on capillary walls, denoted by a white oval, and white lines indicate areas of pericyte site and non-pericyte measurements. The C57BL/6J (C57) mouse *in situ* vasa recta have a comparable appearance to the rodent vasa recta. Scale bar measures 10 μm. Vessel diameter was measured during superfusion of tissue slices with PSS, C5a and a subsequent wash with PSS. The line graph (Aii) shows a representative trace of these measured changes in rodent vessel diameter at pericyte (black line) and non-pericyte sites (grey line), showing vessel diameter reduced during exposure of C5a (grey box) at pericyte sites with little change at the non-pericyte site. B) Graph shows percentage change in rodent vasa recta diameter at pericyte (black) and non-pericyte sites (grey) in response to C5a, IL-33, TNF-α, and IL-18. TNF-α and C5a were used on C57 tissue as they were more potent components, with IL-1β included as a non-responsive control. Ci) shows a representative murine trace as described in (Aii). Cii) Graph shows percentage change in vasa recta diameter at pericyte sites in SD (black) and the C57 (grey) tissue slices in response to C5a, TNF-α, and IL-1β. Significance was calculated using a two-tailed paired Student’s *t*-test for comparisons between pericyte and non-pericyte sites, and a two-tailed unpaired *t*-test was used for comparisons between SD and C57. For comparison between the immune components a one-way ANOVA and post hoc Tukey’s test were used. #P<0.05 between component values; ***P < 0.001 for all other comparisons. Black lines and error bars show means ± SEM; n ≥ 3 animals.

The C57BL/6J mouse is known to be relatively resistant to vasoconstrictors compared to the SD rat (20). Given that C5a and TNF-α elicited a significantly greater magnitude of constriction in rat tissue than that elicited by IL-18 or IL-33 (P < 0.05), mouse tissue was subsequently only exposed to C5a and TNF-α and IL-1β. Both C5a and TNF-α elicited a pericyte-mediated constriction of vasa recta that was significantly greater at pericyte sites (7.4 ± 1.4% and 8.6 ± 0.6% respectively) than non-pericyte sites (2.7 ± 1.0; n = 5; and 2.5 ± 1.2 %; n = 3; P < 0.05). IL-1β failed to elicit a significantly greater change in capillary diameter (4.9 ± 1.6%) at pericyte sites compared to non-pericyte sites (2.8 ± 0.9; n = 4; P > 0.05). Interestingly, the temporal aspect of the contractile responses to C5a and TNF-α appeared to be different in the two species (Fig. 1Aii and Fig. 1Cii for exemplary C5a traces in rats and mice respectively). Murine contractile responses to C5a were of significantly lower magnitude than those recorded in the rat (Fig. 1Ci, p<0.001), whilst TNF-α and iL-1β were not significantly different (Fig 1Cii). Only the rat slice model was used to further investigate if cytokine-evoked vasoconstriction involves angiotensin II type 1 receptors (AT_1_-R), as mouse vasa recta diameter did not return to baseline following removal of either C5a or TNF-α within the experimental timeframe.

### Acute innate immune component-mediated vasoconstriction is AT_1_ receptor dependent

To determine whether the vasoconstriction evoked by C5a, IL-33, IL-18 and TNF-α involved AT^1^-R’s, live kidney slices were superfused with the innate immune components alone and in the presence of the AT_1_ receptor antagonist losartan (100 nM). Application of C5a, IL-33, IL-18 and TNF-α alone prior to inclusion of losartan in the superfusate induced a pericyte-mediated constriction (13.9% ± 0.9%, 9.6% ± 1.1%, 8.9% ± 0.6%, 10.3% ± 0.9%; Fig. 2) of vasa recta. The inclusion of losartan in the superfusate with each immune component, attenuated the pericyte-mediated constriction of vasa recta in response to C5a, IL-33, IL-18 and TNF-α by 88.5%, 95.8%, 96.6% and 71.8% respectively (n = 3, p < 0.05, Fig. 2). Losartan-evoked attenuation of pericyte-mediated constriction was reversible, reapplication of TNF-α in the absence of losartan resulted in a second pericyte-mediated vasoconstriction (14.3% ± 2.3%; n = 4; ***P < 0.001; Fig. 2A) that was not significantly different (P > 0.05) to the first response recorded (10.3% ± 0.6%; Fig. 2A). The losartan-induced attenuation of pericyte-mediated constriction was therefore not thought to be due to desensitisation of AT_1_ receptors. No significant changes in vessel diameter were observed at non-pericyte sites throughout any of these experiments (data not shown).

**Figure 2.**
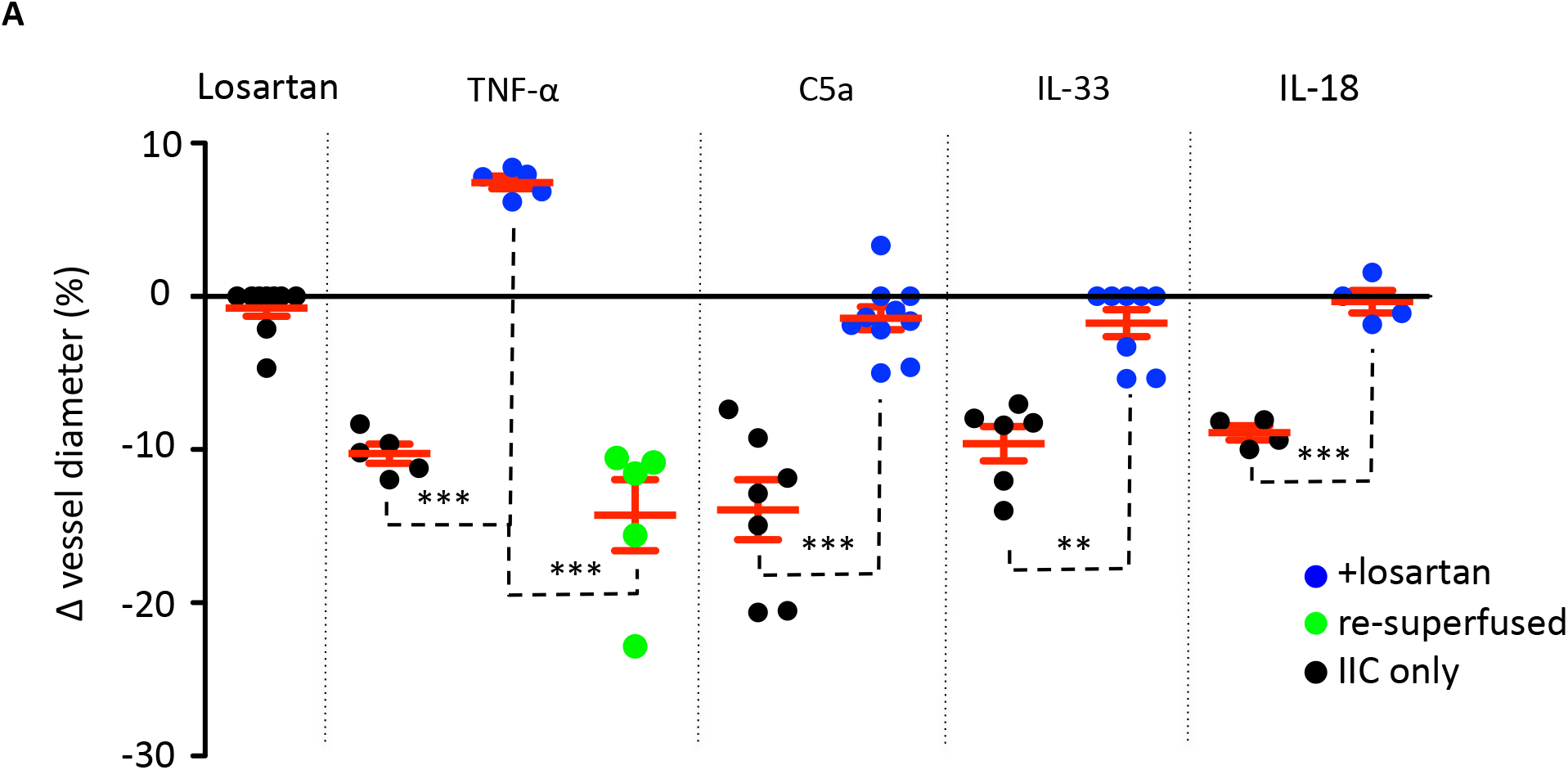
Losartan reversibly attenuates TNF-α-, C5a-, IL-33-, and IL-18-mediated constriction of rodent vasa recta. Graph shows percentage changes in rodent vasa recta diameter at sites in response to superfusion of tissue with innate immune components (IIC only; black spheres), followed by innate immune components combined with the AT_1_ antagonist losartan (blue spheres). When tissue was superfused with TNF-α, C5a, IL-33, and IL-18 alone, a significant constriction of vasa recta was measured at pericyte sites compared to non-pericyte sites. The addition of losartan significantly attenuated the pericyte mediated constriction evoked by TNF-α, C5a, IL-33 and IL-18. In TNF-α experiments tissue was subsequently re-superfused with TNF-α alone (green spheres). Re-superfusion of tissue with TNF-α evoked a significant constriction of vasa recta at pericyte sites, indicating that the effects of losartan were reversible. No significant change in vessel diameter was observed at non-pericyte sites. Superfusion of tissue with losartan alone had no significant effect on vessel diameter at pericyte or non-pericyte sites. Significance was calculated using a two-tailed unpaired Student’s *t*-test. **P < 0.01, ***P < 0.001. Values are means ± SEM; n ≥ 5 animals.

To determine whether endogenous Ang-II activity may be contributing to pericyte-mediated vasoconstriction independently of C5a, IL-33 and TNF-α, kidney slices were superfused with losartan alone and then co-applied with immune components. Losartan alone caused no significant change in vessel diameter at pericyte sites (Fig. 2) compared to non-pericyte sites (P > 0.05; data not shown). Subsequent inclusion of innate immune components, C5a, IL-33, IL-18 and TNF-α in the superfusate with losartan, failed to elicit and further change in vessel diameter at pericyte or non-pericyte sites (P > 0.05; data only shown for TNF-α; Fig. 2).

### Prolonged exposure of kidney tissue to inflammatory mediators alters pericyte density, morphology and vessel diameter

Having investigated the effect of acute exposure of kidney tissue to a series of innate immune components, we sought to further investigate the impact of exposing live kidney tissue to the same components for a period of 4 hours. Following fixation of this tissue and fluorescence imaging of immunohistochemically labelled percytes and vasa recta capillaries, we measured the denisity of NG2^+^ pericyte cells along vasa recta capillaries (in 100 μm^2^ of tissue), pericyte process length (around vasa recta), pericyte cell body length and width, and vessel diameter (at pericyte site and non-pericyte sites). Fluorescence images of fixed tissue slices exposed to immune components and PSS alone were acquired (Fig. 3A-C) and analysed off-line. Exposure of rodent tissue to IL-33, C5a, IL-18 and TNF-α (4 hours, compared to 4 hour treatment with PSS alone), prior to fixation, evoked a significant decrease in pericyte density (Fig. 3D), vessel diameter at pericyte sites (Fig. 3E), and circumferential pericyte process length (Fig. 3F) compared to those measured in tissue exposed to PSS only (see Table 1 for values, n = 3). There was no significant difference in vessel diameter (Fig. 3E) at non-pericyte sites in response to innate immune components compared to PSS control experiments (see Table 1 for values, n = 3). There was no significant change in pericyte cell body width in response to IL-33, IL-18 or TNF-α whereas pericyte cell width decreased in response to C5a compared to PSS control experiments (Fig. 3F, see Table 1 for values, n = 3). There was also a significant increase in pericyte cell body height recorded in tissue exposed to C5a, IL-33, IL-18 and TNF-α, compared to that measured for pericytes exposed to the PSS only (Fig. 3F, see Table 1 for values, n = 3).

**Table 1.** Summary table of anti-NG2 immunohistochemistry experiment data. Table shows the measurements for pericyte density per 100 μm^2^ (Pericyte no.), pericyte morphology, and vessel diameter in slices exposed to innate immune components and PSS and corresponding significance values (PSS vs innate immune component). Values are means ± SEM; n = 3 animals. A two-tailed paired Student’s t-test was used to calculate significance in the C57BL/6J mouse. Significance in the Sprague-Dawley rat was calculated using a one-way ANOVA and post hoc Dunnet test. *P < 0.05, **P < 0.01, ***P < 0.001.

| Innate immune component | Pericyte no. | Cell body width ($\mu\text{m}$) | Cell body height ($\mu\text{m}$) | Process length across vessel ($\mu\text{m}$) | Vessel diameter at pericyte sites ($\mu\text{m}$) | Vessel diameter at non-pericyte sites ($\mu\text{m}$) | Fold increase vs acute superfusion |
| --- | --- | --- | --- | --- | --- | --- | --- |
| Sprague-Dawley Rat |  |  |  |  |  |  |  |
| **PSS** | 19.0 ± 1.3 | 9.3 ± 0.3 | 3.5 ± 0.2 | 8.6 ± 0.4 | 8.2 ± 0.3 | 10.2 ± 0.5 | N/A |
| **C5a** | 9.0 ± 0.7<br>***P < 0.001 | 7.6 ± 0.3<br>***P < 0.001 | 5.2 ± 0.2<br>***P < 0.001 | 5.8 ± 0.5<br>***P < 0.001 | 5.6 ± 0.4<br>**P > 0.01 | 9.0 ± 0.5<br>P > 0.05 | (15.2 vs 32%)<br>approximately double |
| **IL-33** | 12.0 ± 1.0<br>***P < 0.001 | 8.5 ± 0.3<br>P > 0.05 | 4.8 ± 0.1<br>***P < 0.001 | 5.8 ± 0.4<br>***P < 0.001 | 6.7 ± 0.3<br>**P > 0.01 | 10.6 ± 0.4<br>P > 0.05 | (9.21 vs 19%)<br>approximately double |
| **TNF-α** | 12.8 ± 0.4<br>***P < 0.001 | 8.6 ± 0.2<br>P > 0.05 | 5.1 ± 0.1<br>***P < 0.001 | 7.3 ± 0.2<br>**P > 0.01 | 5.0 ± 0.2<br>***P < 0.001 | 9.0 ± 0.3<br>P > 0.05 | (11.7 vs 39%)<br>approximately 4-fold |
| **IL-18** | 12.3 ± 0.8<br>***P < 0.001 | 8.6 ± 0.2<br>P > 0.05 | 4.9 ± 0.2<br>***P < 0.001 | 6.9 ± 0.3<br>**P > 0.01 | 6.3 ± 0.2<br>***P < 0.001 | 9.5 ± 0.2<br>P > 0.05 | (9.8 vs 23%)<br>approximately 2.5-fold |
| C57BL/6J Mouse |  |  |  |  |  |  |  |
| **PSS** | 12.6 ± 1.0 | 7.5 ± 0.4 | 4.5 ± 0.2 | 6.8 ± 0.5 | 6.3 ± 0.3 | 8.1 ± 0.4 | N/A |
| **TNF-α** | 8.8 ± 0.9<br>*P < 0.05 | 7.4 ± 0.5<br>P > 0.05 | 4.3 ± 0.3<br>P > 0.05 | 5.1 ± 0.4<br>*P < 0.05 | 5.1 ± 0.3<br>*P < 0.05 | 6.9 ± 0.4<br>*P < 0.05 | (8.6% vs 17%)<br>approximately double |

**Figure 3.**
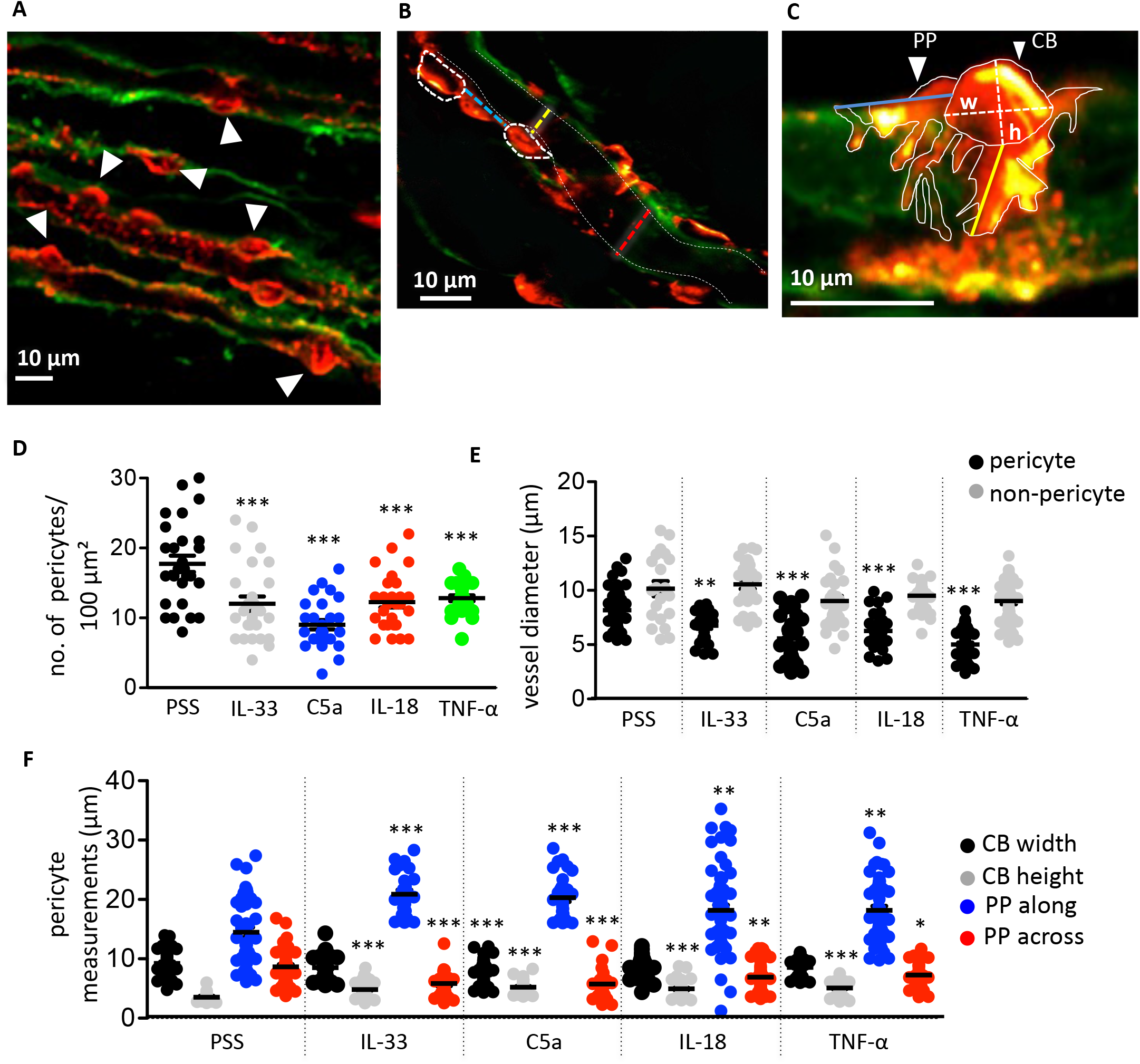
C5a, IL-33, IL-18 and TNF-α evoked changes in pericyte density, physiology, and vessel diameter in rat tissue. A-C) Images show typical fields of view of Sprague-Dawley rodent vasculature labeled with IB_4_ (green) and pericytes labeled with anti-NG2 (red). (A) Image shows pericyte density (white arrowheads indicating pericytes) in an area of 100 μm^2^. (B) shows pericyte distribution along an individual vasa recta (white dashed ovals outline pericyte cell bodies (CB)) with the distance between indicated by the blue dashed line. Yellow and red dashed lines indicate where vasa recta diameter was measured at a pericyte and non-pericyte site respectively. White dashed lines indicate the capillary walls. C) Image shows pericyte CB and pericyte processes (PP) outlined with solid white lines. Where the CB height (h) and width (w) measurements taken are denoted by dashed white lines. Longitudinal (PP along) and circumferential (PP across) process measurements for individual pericyte soma were also taken, indicated by the blue and yellow lines respectively. Graphs show the number of pericytes within an area of 100 μm^2^ (D), vessel diameter at pericyte and non-pericyte sites (E), and (F) shows both CB width and height, and PP length along and across vessels. All stimuli evoked a significant decrease in pericyte density in an area of 100 μm^2^, compared to PSS time matched control group. A significant decrease in process length across vessels and an increase in process length around vessels was also recorded in response to all stimuli compared to PSS control group. Vasa recta diameter was also significantly reduced at pericyte-site sites with no significant change measured at non-pericyte sites. Exposure of tissue to all stimuli evoked a significant increase in CB height compared to PSS time matched control group, however, there was no significant difference between CB width in response to the innate immune components apart from C5a that showed a significant reduction in CB width compared to PSS control group. Data in (D) is count data and as such significance was calculated using a negative binomial regression with the form *Density ∼ treatment + (1* | *animal)*. Significance for all other data here was calculated using a one-way ANOVA with post hoc Dunnet test. *P < 0.05, **P < 0.01, ***P < 0.001. Black lines and error bars show means ± SEM; n = 3 animals.

Given that there was no significant difference in the TNF-α mediated constriction of vasa recta in mouse tissue when compared to that in rat (unlike C5a), mouse tissue was exposed to TNF-α only in this set of experiments. In murine tissue, the basal pericyte density was lower than in rats, and the vasa recta capillaries are narrower (Table 1; Fig. 3D and Fig. 4A). Exposure of murine tissue to TNF-α, prior to fixation, evoked a significantly greater decrease in pericyte density (P < 0.05; Fig. 4A), vasa recta diameter (P < 0.05; Fig. 4B) and circumferential process length (P < 0.05; Fig. 4C) compred to that measured in mouse tissue exposed to PSS alone (see Table 1 for values, n = 3). No significant changes in pericyte cell body width or height were detected in tissue treated with TNF-α when compared to tissue treated with PSS alone (Fig. 4C).

**Figure 4.**
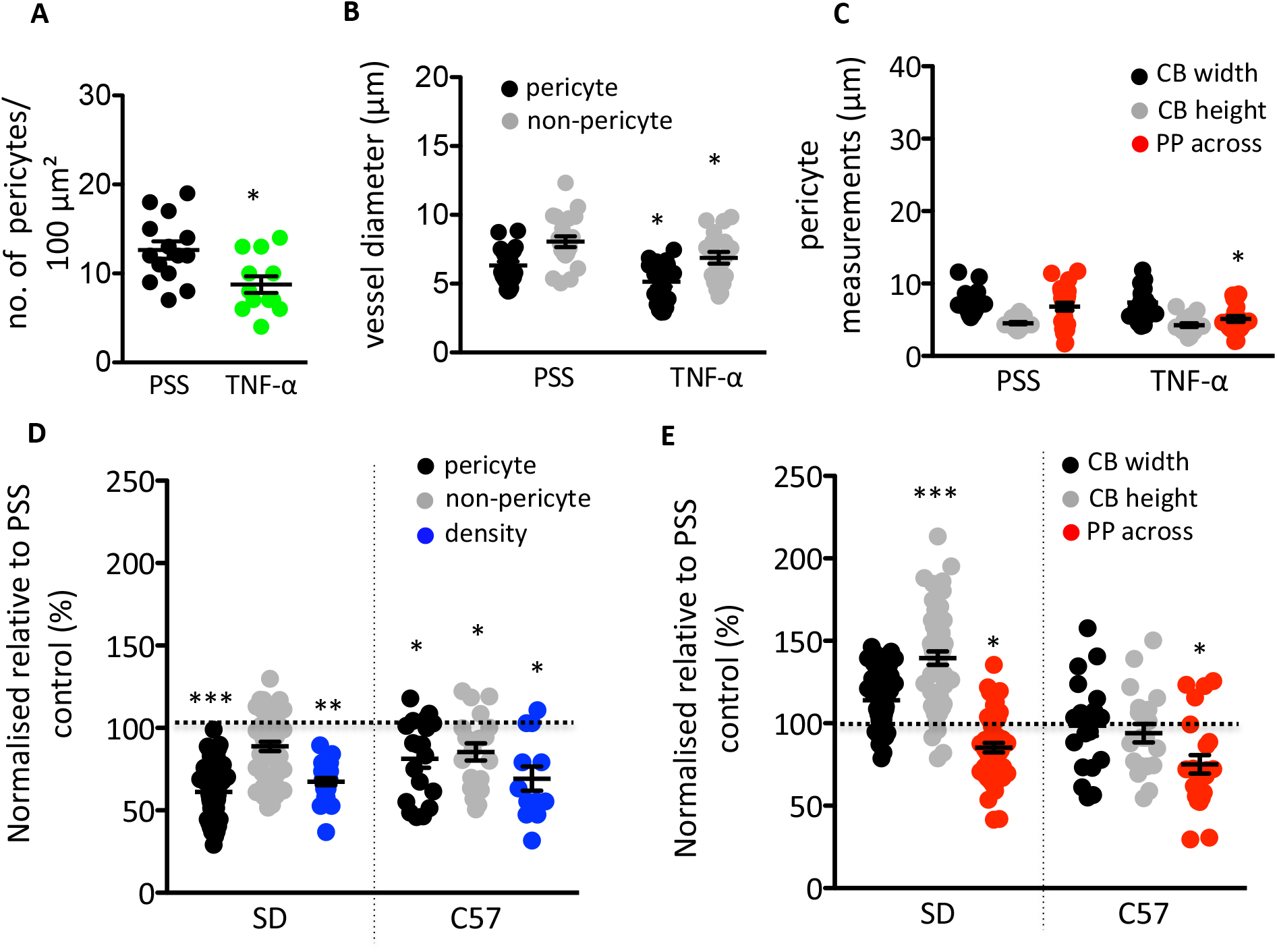
TNF-α stimulation of murine tissues evokes comparable changes in pericyte density, pericyte processes, and vasa recta diameter to those in measured the rat. Graphs show the results of TNF-α stimulation of C57BL/6J (C57) tissue comparative to the PSS time-matched control. A) shows the number of pericytes within an area of 100 μm^2^, B) the vessel diameter at pericyte and non-pericyte sites, and C) circumferential pericyte processes (PP across), and pericyte cell body (CB) height and width following 4-hours of either TNF-α or control PSS stimulation. TNF-α evoked a significant decrease in pericyte density per 100 μm^2^, comparative to the PSS time matched control group. A significant decrease in process length across vessels, and vasa recta diameter at both pericyte-site sites and non-pericyte sites was measured after incubation with TNF-α. TNF-α stimulation had no effect of murine pericyte CB characteristics. (D) Graph shows PSS-normalised values for vasa recta diameter at pericyte and non-pericyte sites, and pericyte density (density) in Sprague-Dawley (SD) rats and C57 upon exposure to TNF-α, with (E) showing PSS-normalised data for all other measurements taken (CB width and height, and PP across). The PSS average for each metric was set as “100%” (black dashed line). What can be seen are some consistent cross-species effects after TNF-α stimulation, with changes in CB morphology unique to the rat. A two-tailed paired Student’s *t*-test was used to calculate significance in the C57.Significance in the SD was calculated using a one-way ANOVA and post hoc Dunnet test. All statistical values are for PSS vs TNF-α; *P < 0.05, **P < 0.01, ***P < 0.001. Black lines and error bars show means ± SEM; n = 3 animals in C57.

When comparing the normalized TNF-α evoked changes in mouse and rat tissue, it was noted that the relative change in pericyte density (Fig. 4D), vasa recta diameter (Fig. 4D), and circumferential processe length (Fig. 4E) after exposure to TNF-α were not significantly different between species.

## Discussion

In the present study we have demonstrated that (i) innate immune components can elicit a pericyte-mediated constriction of vasa recta, (ii) they elicit this acute vasoconstriction via activity at Ang-II receptor type 1 (AT_1_-R), and (iii) sustained exposure to inflammatory mediators led to a reduction in number and morphological changes to NG2^+^ pericyte soma and processes. Taken together, we have shown inflammatory mediators act at renal pericytes in the medulla to dysregulate vasa recta diameter and thus are potentially involved in the decline in MBF and may induce cellular events that are associated with AKI.

Circulating immune cells release pro-inflammatory mediators as they bind to the endothelium (21). Several studies, covering a range of species and vascular beds, have shown inflammatory mediators can be directly and indirectly vasoactive (8,22,23) Direct vasoactivity involves action at cytokine receptors on endothelial, vascular smooth muscle cells (vSMC)(8,24), and pericytes (25). Indirect vasoactivity involves the downstream release of vasodilatory mediators like NO and PGI_2_ by upregulating COX-2 and iNOS, as well as stimulating endothelial production of vasoconstrictive mediators including ROS, Ang-II and ET-1 (8,22,23). Our imaging of live rodent and murine kidney tissue enabled us to show in real-time how IRI-associated complement proteins (C5a) and cytokines (IL-33, IL-18 and TNF-α) act at vasa recta pericytes to constrict DVR in an acute setting (Fig 1). Further still, our study is the first to show that inflammatory mediators elicit an acute pericyte-mediated constriction in an Ang-II type 1 receptors (AT_1_-R)-dependent manner (Fig 2) in the rat. There is an interconnected relationship between inflammatory mediators and Ang-II (9), with AT_1_-R blocker losartan impairing post-ischaemic cytokine production and limiting injury (26,27). Whilst not the most significant pathway involved in IRI induced changes to medullary blood flow, the involevement of AT_1_-R activation has been implicated in no-reflow following renal IRI (19). In experiements shown here, superfusion with losartan alone elicited no change in vessel diameter, whilst superfusion of losartan together with TNF-α, C5a, IL-18, or IL-33 significantly attenuated the innate immune component-induced vasoconstriction (Fig. 2). This data set indicates that innate immune component-mediated constriction involves activation of AT_1_-R by Ang-II that is not present in un-stimulated tissue (Fig 2). This data agrees with other studies where C3a induces Ang-II production in vSMC from spontaneously hypertensive rats (SHR) (28) and both TNF-α (23) and C5a (29) are linked with Ang-II in murine models of Ang-II-induced hypertension, CKD, and cardiac injury.

The vasoctivity of innate immune components involving AT_1_-R could explain vascular-bed specific differences in cytokine behaviour, and the species difference in magnitude of pericyte-mediated constriction (Fig. 1Cii). Upon exposure to TNF-α, a vasodilation is measured in the rodent cremaster muscle (24) whilst it constricts the rodent and murine renal vasculature (22,23). If we relate this vasoactivity to AT-R distribution, the rodent cremaster muscle expresses more dilatory AT_2_-R (30) than rodent kidneys (31), and offers an explanation for the observed differences. An AT-R dependent difference in Ang-II contractility has also been shown in the mesentery of aged mice where reduction in AT_2_-R approximately doubles the contractility of Ang-II comparative to young mice (32). Moreover, the SD rodent kidney has approximately double the AT_1_-R density of the C57BL/6J mouse (33), and thus a greater magnitude of pericyte-mediated contractility. Given the fundamental role of the renin-angiotensin system in kidney function it is perhaps unsurprising that there might be a role for AT_1_-R in the immune-component elicited pericyte-mediated constriction of DVR, however only performing losartan experiments in rat tissue means these finding may not generalise as described here and not repeating these experiments in mice is a noted limitation of this study.

Conflicting somewhat with other work highlighting roles for C3a and IL-1β in the pathogenesis of IRI-AKI (34,35) these immune components failed to elicit acute pericyte-mediated changes in DVR diameter (Fig. 1), yet there are other ways they may contribute to heamodynamic dysregulation. In the SHR, IL-1β potentiates the vasoactive response of phenylephrine, ET-1, and Ang-II in the mesentery, cremaster and heart (24,36–38) opposed to direct vasoactivity. Our acute experimental window may also be insufficient to induce indirect vasomotor activity; the cytokine upregulation induced by C5a and C3a is time-dependent (39). Another consideration is non-vasoactive actions that cause injury and may lead to impaired MBF. Antagonism of IL-1 receptors has been shown to alter lymphocyte and macrophage infiltration, and IL-1α/β knock out mice exhibit reduced acute tubular necrosis between 24 and 48 hours after ischemia (34). This immune cell infiltrate occludes vasa recta (3,4,40–42) and could impede blood flow independently of the actions of pericytes. So, whilst C3a and IL-1β contribute to kidney injury (34,35), these mechanisms appear independent from acute pericyte-mediated MBF dysregulation and is beyond the scope of this study.

In longer term experiments, where tissue was exposed to innate immune components for 4 hours, vessels remain strangulated by pericytes in response to C5a, IL-33, IL-18 (Fig. 3E) and TNF-α (Fig. 3E and 5B). In these incubations the vessel constriction was not only sustained but approximately double that of the acute experiments (Table 1). Previously, we, and others, have argued that pericyte-mediated changes in DVR diameter are likely to underpin localized changes in blood flow, and dysregulation of MBF via pericyte cells is likely to underpin many renal pathologies (14,43–46). Whilst we cannot say for certain constriction was sustained from the start of incubation, four hours of reduced blood flow, due to pericyte strangulation, is a substantial amount of time for tissue to be under perfused and likely to cause ischaemic and hypoxic conditions. Recently it has been shown that NG2^+^ pericytes mediate the medullary no-reflow in renal ischaemia (19), we further propose that that renal injury-associated innate-immune components tested here, specifically C5a, IL-33, IL-18 and TNF-α, might also play a role in the sustained dysregulation of MBF that occurs during ischemia-reperfusion.

Experiments presented here show that C5a, IL-33, IL-18 and TNF-α evoke a decrease in pericyte density and elicit changes in pericyte morphology after only 4 hours of exposure (Fig. 3 and 5). Firstly, we reveal that innate immune components C5a, IL-33, TNF-α and IL-18 evoke a decrease in pericyte density across the medulla (Fig. 3D and 5A). In porcine IRI, inhibiting C5a ameliorates the significant reduction in NG2^+^ pericyte number and capillary constriction at 24 hours post-reperfusion (47), and in SHR this loss of NG2^+^ pericytes is associated with increased injury following ischaemia (48). However, we are the first to further quantify changes in pericyte morphology in response to innate immune components. We have revealed that C5a, IL-33, TNF-α and IL-18 induce changes in circumferential pericyte process length (decrease) vasa recta capillaries (Fig. 3F and 5C), whilst only increasing cell body height in rats (Fig. 3F, 5E). These findings are supported by a previous study where injection of TNF-a and IL-1β induced a loose and non-confluent coverage of pericytes along postcapillary venules in murine cremaster muscle and ear skin (49). TNF-α stimulation encourages an elongated migratory morphology of pericytes (50) and a retraction of pericyte processes (51). Whilst this relaxation of pericyte processes is necessary for leukocyte extravasation (52), this process could initiate the pericyte-myofibroblast transition as this pericyte relaxation (52), leukocyte migration (53), and fibrotic development (54) are Rho-A/ROCK dependent, though further work is needed to elucidate this mechanism. Freitas and Attwell (19) further demonstrated how Rho-A/ROCK inihibition restores MBF fastest following IRI, and inihibits the no-reflow phenomenon specifically by activity at NG2^+^ pericytes. Pericytes provide stability to the capillaries on which they reside (14), and their loss induces capillary rarefaction (18) which culminates in a loss of up to 50% of the vasa recta capillaries in rodents 4-weeks after ischaemia (6,55). Our data here, and others (47,48), demonstrating an early reduction of NG2^+^ pericytes is possibly linked to the vessel instability and rarefaction observed renal disease (14,18), with morphological changes potentially suggesting a fibrotic phenotype of these lost pericytes (14–17).

## Conclusions

In conclusion, we have provided novel evidence demonstrating inflammatory mediators can evoke acute pericyte-mediated changes in vasa recta diameter, as well as changes in NG2^+^ pericyte density and morphology. These effects could underpin the sustained dysregulation of MBF in IRI, leading to hypoxic cellular and tissue damage and subsequent renal dysfunction. All of these mechanisms have been linked with AKI and the transition to CKD, and possibly underlie early cellular events that lead to pericyte detachment and differentiation into myofibroblasts. Whilst further work is needed, delineating these mechanisms could highlight potential therapeutic targets to prevent the onset of AKI following IRI.

## References

1. Ponce D, Balbi A. Acute kidney injury: risk factors and management challenges in developing countries. Int J Nephrol Renovasc Dis. 2016;9:193–200.

2. Imig JD, Ryan MJ. Immune and inflammatory role in renal disease. Compr Physiol. 2013;3(2):957–76.

3. Kielar ML, John R, Bennett M, Richardson JA, Shelton JM, Chen L, et al. Maladaptive role of IL-6 in ischemic acute renal failure. J Am Soc Nephrol. 2005 Nov 1;16(11):3315–25.

4. Basile D, Anderson M, Sutton T. Pathophysiology of Acute Kidney Injury. Compr Physiol [Internet]. 2012;2(2):1303–53. Available from: http://www.ncbi.nlm.nih.gov/pmc/articles/PMC3919808/?report=reader

5. Ray SC, Mason J, O’Connor PM. Ischemic Renal Injury: Can Renal Anatomy and Associated Vascular Congestion Explain Why the Medulla and Not the Cortex Is Where the Trouble Starts? Seminars in Nephrology [Internet]. 2019;39(6):520–9. Available from: https://doi.org/10.1016/j.semnephrol.2019.10.002

6. Regner KR, Roman RJ. Role of medullary blood flow in the pathogenesis of renal ischemia–reperfusion injury. Current Opinion in Nephrology and Hypertension. 2012 Jan;21(1):33–8.

7. Pallone TL, Edwards A, Mattson DL. Renal medullary circulation. Compr Physiol. 2012;2(1):97–140.

8. Vila E, Salaices M. Cytokines and vascular reactivity in resistance arteries. American Journal of Physiology - Heart and Circulatory Physiology. 2005;288(3 57-3):1016–21.

9. Liu Z, Huang XR, Lan HY. Smad3 mediates ANG II-induced hypertensive kidney disease in mice. AJP: Renal Physiology. 2012 Apr 15;302(8):F986–97.

10. Bonventre J V., Zuk A. Ischemic acute renal failure: An inflammatory disease? Kidney International. 2004 Aug;66(2):480–5.

11. Crawford C, Kennedy-Lydon T, Sprott C, Desai T, Sawbridge L, Munday J, et al. An intact kidney slice model to investigate vasa recta properties and function in situ. Nephron - Physiology. 2012;120(3).

12. Crawford C, Kennedy-Lydon TM, Callaghan H, Sprott C, Simmons RL, Sawbridge L, et al. Extracellular nucleotides affect pericyte-mediated regulation of rat in situ vasa recta diameter. Acta Physiol (Oxf). 2011;202(3):241–51.

13. Cowley Jr AWC, O’connor PM. MEDULLARY THICK ASCENDING LIMB BUFFER VASOCONSTRICTION OF RENAL OUTER-MEDULLARY VASA RECTA IN SALT-RESISTANT BUT NOT SALT-SENSITIVE RATS. Hypertension. 2013;60(4):965–72.

14. Lemos DR, Marsh G, Huang A, Campanholle G, Aburatani T, Dang L, et al. Maintenance of vascular integrity by pericytes is essential for normal kidney function. American Journal of Physiology-Renal Physiology. 2016 Dec;311(6):F1230–42.

15. Kramann R, Schneider RK, DiRocco DP, Machado F, Fleig S, Bondzie PA, et al. Perivascular Gli1+ progenitors are key contributors to injury-induced organ fibrosis. Cell Stem Cell. 2015 Jan 8;16(1):51–66.

16. Lin SL, Kisseleva T, Brenner D a, Duffield JS. Pericytes and Perivascular Fibroblasts Are the Primary Source of Collagen-Producing Cells in Obstructive Fibrosis of the Kidney. The American Journal of Pathology. 2008;173(6):1617–27.

17. Humphreys BD, Lin SL, Kobayashi A, Hudson TE, Nowlin BT, Bonventre J V, et al. Fate tracing reveals the pericyte and not epithelial origin of myofibroblasts in kidney fibrosis. Am J Pathol. 2010;176(1):85–97.

18. Kramann R, Wongboonsin J, Chang-Panesso M, Machado FG, Humphreys BD. Gli1 ^+^ Pericyte Loss Induces Capillary Rarefaction and Proximal Tubular Injury. Journal of the American Society of Nephrology. 2017;28(3):776–84.

19. Freitas F, Attwell D. Pericyte-mediated constriction of renal capillaries evokes no-reflow and kidney injury following ischaemia. Elife. 2022;11:1–26.

20. Russell A, Watts S. Vascular reactivity of isolated thoracic aorta of the C57BL/6J mouse. J Pharmacol Exp Ther [Internet]. 2000;294(2):598–604. Available from: http://www.ncbi.nlm.nih.gov/pubmed/10900237

21. Bonventre J, Yang L. Cellular pathophysiology of ischemic acute kidney injury. J Clin Invest. 2011;121(11):4210–21.

22. Shahid M, Francis J, Majid D. Tumor necrosis factor-α induces renal vasoconstriction as well as natriuresis in mice. American Journal of Physiology - Renal Physiology. 2008;295(6):F1836–44.

23. Zhang J, Patel MB, Griffiths R, Mao A, Song Y soo, Karlovich NS, et al. Tumor necrosis factor-α produced in the kidney contributes to angiotensin II-dependent hypertension. Hypertension. 2014 Dec;64(6):1275–81.

24. Baudry N, Rasetti C, Vicaut E. Differences between cytokine effects in the microcirculation of the rat. Am J Physiol. 1996 Sep;271(3 Pt 2):H1186–92.

25. Kerkar S, Williams M, Blocksom JM, Wilson RF, Tyburski JG, Steffes CP. TNF-α and IL-1β Increase Pericyte/Endothelial Cell Co-Culture Permeability. Journal of Surgical Research. 2006;132(1):40–5.

26. Molinas SM, Cortés-González C, González-Bobadilla Y, Monasterolo LA, Cruz C, Elías MM, et al. Effects of losartan pretreatment in an experimental model of ischemic acute kidney injury. Nephron - Experimental Nephrology. 2009;112(1).

27. Gabriele LG, Morandini AC, Dionísio TJ, Santos CF. Angiotensin II Type 1 Receptor Knockdown Impairs Interleukin-1β-Induced Cytokines in Human Periodontal Fibroblasts. J Periodontol. 2017;88(1):e1–11.

28. Han Y, Fukuda N, Ueno T, Endo M, Ikeda K, Xueli Z, et al. Role of complement 3a in the synthetic phenotype and angiotensin II-production in vascular smooth muscle cells from spontaneously hypertensive rats. Am J Hypertens. 2012 Mar;25(3):284–9.

29. Zhang C, Li Y, Wang C, Wu Y, Cui W, Miwa T, et al. Complement 5a receptor mediates angiotensin II-induced cardiac inflammation and remodeling. Arterioscler Thromb Vasc Biol. 2014 Jun;34(6):1240–8.

30. Hong K, Zhao G, Hong Z, Sun Z, Yang Y, Clifford PS, et al. Mechanical activation of angiotensin II type 1 receptors causes actin remodelling and myogenic responsiveness in skeletal muscle arterioles. J Physiol. 2016;594(23):7027–47.

31. Ruan X, Wagner C, Chatziantoniou C, Kurtz A, Arendshorst WJ. Regulation of angiotensin II receptor AT1 subtypes in renal afferent arterioles during chronic changes in sodium diet. Journal of Clinical Investigation. 1997;99(5):1072–81.

32. Dinh QN, Drummond GR, Kemp-Harper BK, Diep H, De Silva TM, Kim HA, et al. Pressor response to angiotensin II is enhanced in aged mice and associated with inflammation, vasoconstriction and oxidative stress. Aging. 2017;9(6):1595–606.

33. Cassis LA, Huang J, Gong MC, Daugherty A. Role of metabolism and receptor responsiveness in the attenuated responses to Angiotensin II in mice compared to rats. Regulatory Peptides. 2004;117(2):107–16.

34. Rusai K, Huang H, Sayed N, Strobl M, Roos M, Schmaderer C, et al. Administration of interleukin-1 receptor antagonist ameliorates renal ischemia-reperfusion injury. Transplant International. 2008 Jun;21(6):572–80.

35. Peng Q, Li K, Smyth LA, Xing G, Wang N, Meader L, et al. C3a and C5a promote renal ischemia-reperfusion injury. J Am Soc Nephrol. 2012 Sep;23(9):1474–85.

36. Dorrance AM. Interleukin 1-beta (IL-1beta) enhances contractile responses in endothelium-denuded aorta from hypertensive, but not normotensive, rats. Vascul Pharmacol. 2007;47(2–3):160–5.

37. De Salvatore G, De Salvia MA, Piepoli AL, Natale L, Porro C, Nacci C, et al. Effects of in vivo treatment with interleukins 1beta and 6 on rat mesenteric vascular bed reactivity. Auton Autacoid Pharmacol. 2003 Apr;23(2):125–31.

38. Vicaut E, Rasetti C, Baudry N. Effects of tumor necrosis factor and interleukin-1 on the constriction induced by angiotensin II in rat aorta. Journal of Applied Physiology. 1996 Jun 1;80(6):1891–7.

39. Monsinjon T, Gasque P, Chan P, Ischenko A, Brady JJ, Fontaine MC. Regulation by complement C3a and C5a anaphylatoxins of cytokine production in human umbilical vein endothelial cells. FASEB J. 2003;17(9):1003–14.

40. Naruse T, Yuzawa Y, Akahori T, Mizuno M, Maruyama S, Kannagi R, et al. P-selectin-dependent macrophage migration into the tubulointerstitium in unilateral ureteral obstruction. Kidney International. 2002;62(1):94–105.

41. Kelly KJ, Williams WW, Colvin RB, Meehan SM, Springer TA, Gutiérrez-Ramos JC, et al. Intercellular adhesion molecule-1-deficient mice are protected against ischemic renal injury. Journal of Clinical Investigation. 1996;97(4):1056–63.

42. Ysebaert DK, De Greef KE, De Beuf A, Van Rompay ANR, Vercauteren S, Persy VP, et al. T cells as mediators in renal ischemia/reperfusion injury. Kidney International. 2004 Aug;66(2):491–6.

43. Peppiatt-Wildman CM. The evolving role of renal pericytes. Current Opinion in Nephrology and Hypertension. 2013 Jan;22(1):10–6.

44. Kennedy-Lydon TM, Crawford C, Wildman SSP, Peppiatt-Wildman CM. Renal pericytes: Regulators of medullary blood flow. Acta Physiologica. 2013;207(2):212–25.

45. Kennedy-Lydon T, Crawford C, Wildman SS, Peppiatt-Wildman CM. Nonsteroidal anti-inflammatory drugs alter vasa recta diameter via pericytes. American Journal of Physiology - Renal Physiology. 2015;309(7):F648–57.

46. Xavier S, Sahu RK, Landes SG, Yu J, Taylor RP, Ayyadevara S, et al. Pericyte and immune cells contribute to complement activation in tubulointerstitial fibrosis. American Journal of Physiology - Renal Physiology. 2017;ajprenal.00604.2016.

47. Castellano G, Franzin R, Stasi A, Divella C, Sallustio F, Pontrelli P, et al. Complement activation during ischemia/reperfusion injury induces pericyte-to-myofibroblast transdifferentiation regulating peritubular capillary Lumen Reduction Through pERK Signaling. Frontiers in Immunology. 2018;9(MAY):1–17.

48. Crislip GR, O’Connor PM, Wei Q, Sullivan JC, O’Connor PM, Wei Q, et al. Vasa Recta Pericyte Density is Negatively Associated with Vascular Congestion in the Renal Medulla Following Ischemia Reperfusion in Rats. American Journal of Physiology - Renal Physiology [Internet]. 2017;313(5):ajprenal.00261.2017. Available from: http://ajprenal.physiology.org/lookup/doi/10.1152/ajprenal.00261.2017

49. Proebstl D, Voisin MB, Woodfin A, Whiteford J, D’Acquisto F, Jones GE, et al. Pericytes support neutrophil subendothelial cell crawling and breaching of venular walls in vivo. The Journal of Experimental Medicine. 2012;209(6):1219–34.

50. Tigges U, Boroujerdi A, Welser-Alves J V., Milner R. TNF-α promotes cerebral pericyte remodeling in vitro, via a switch from α1 to α2 integrins. Journal of Neuroinflammation. 2013;10:1–15.

51. Dore-Duffy P, Cleary K. Morphology and Properties of Pericytes. In: Methods in molecular biology (Clifton, NJ). 2011. p. 49–68.

52. Wang S, Cao C, Chen Z, Bankaitis V, Tzima E, Sheibani N, et al. Pericytes Regulate Vascular Basement Membrane Remodeling and Govern Neutrophil Extravasation during Inflammation. PLoS ONE. 2012;7(9).

53. Cernuda-Morollón E, Ridley AJ. Rho GTPases and leukocyte adhesion receptor expression and function in endothelial cells. Circulation Research. 2006;98(6):757–67.

54. Baba I, Egi Y, Utsumi H, Kakimoto T, Suzuki K. Inhibitory effects of fasudil on renal interstitial fibrosis induced by unilateral ureteral obstruction. Molecular Medicine Reports. 2015;12(6):8010–20.

55. Basile DP, Donohoe D, Roethe K, Osborn JL. Renal ischemic injury results in permanent damage to peritubular capillaries and influences long-term function. American Journal of Physiology - Renal Physiology. 2001;281(5 50-5):887–99.

